# Mouse SAS-6 is required for centriole formation in embryos and integrity in embryonic stem cells

**DOI:** 10.1101/2022.08.11.503634

**Authors:** Marta Grzonka, Hisham Bazzi

## Abstract

Cell division fidelity is crucial for stem cell propagation and the maintenance of pluripotency. Centrosomes, which organize the mitotic spindle microtubules to ensure proper bipolar cell division, have a core of a pair of centrioles that duplicate once per cell cycle. At the onset of centriole biogenesis, SAS-6 forms a cartwheel structure, which is the precursor for the forming procentrioles. SAS-6 is essential for centriole formation in human cell lines and other organisms. However, the functions of SAS-6 in mouse stem cells remain to be elucidated. Here, we report that *Sas-6*-null mouse embryos lack centrioles, activate the mitotic surveillance cell death pathway and arrest at mid-gestation. In contrast, SAS-6 is not strictly required for centriole formation in mouse embryonic stem cells (mESCs) *in vitro*, but is still important to regulate centriole length, symmetry and ability to template cilia. Remarkably, centrioles appeared after just one day of culture of *Sas-6*-null blastocysts, from which mESCs are normally derived. Finally, the number of cells with centrosomes is drastically decreased upon the exit from a pluripotent state. Collectively, our data suggest a differential requirement for mouse SAS-6 in centriole formation or integrity depending on the cellular context, and highlight the robustness of mESCs in using SAS-6-independent centriole-duplication pathways.

## Introduction

Pluripotent stem cells are characterized by an indefinite capacity of proliferation and self-renewal. To achieve this, they rely on stringent controls of cell division fidelity, primarily through the proper assembly, organization and polarity of the microtubule-based mitotic spindle. These functions are ensured by centrosomes, the major microtubule organizing centers (MTOCs). At the core of centrosomes are the microtubule-based centrioles that are highly conserved in evolution (Bornens, 2012; Nabais *et al*, 2020). During interphase or upon differentiation, centrioles provide the essential template to form cilia (Conduit *et al*, 2015). Cycling cells have two centrioles whose duplication is controlled by the centriole formation machinery. Once per cycle, procentrioles assemble on the existing centrioles and form new daughter centrioles (Nigg & Holland, 2018). The phenotypes of centriole loss vary between organisms and cell types depending on the essential function of centrioles in each context (Marshall, 2007). In humans, various mutations in genes encoding centrosomal and centriolar proteins cause primordial dwarfism and microcephaly (Bond *et al*, 2005; Khan *et al*, 2014). In the mouse, we have shown that centrioles and centrosomes are crucial for embryonic development beyond mid-gestation (Bazzi & Anderson, 2014a, b). Null mutations in mouse *Sas-4*, which is essential for centriole formation, lead to the loss of centrioles and arrest by embryonic day (E) 9 (Bazzi & Anderson, 2014a). The acentriolar cells in the *Sas-4* null embryos activate a p53-, 53BP1- and USP28-dependent cell death pathway, which is known as the mitotic surveillance pathway (Lambrus & Holland, 2017; Xiao *et al*, 2021).

Spindle assembly defective protein 6, SAS-6 (named SASS6), a core protein of the centriole biogenesis pathway (Leidel *et al*, 2005; Nigg & Holland, 2018), forms the major structural component of the cartwheel, which is the precursor for a 9-fold symmetry in centriole assembly and onto which microtubules of the forming procentriole are added (Kitagawa *et al*, 2011). SAS-6 consists of a conserved N-terminal head domain and an intrinsically disordered C-terminal region that flank a conserved coiled-coil domain whose homodimeric association forms a long tail. Hydrophobic interactions between the head domains of the dimers lead to the assembly of the hub, while the tails form the spokes of the cartwheel (Hilbert *et al*, 2013; Kantsadi *et al*, 2022; Kitagawa *et al*., 2011; van Breugel *et al*, 2011; Yoshiba *et al*, 2019). The loss of SAS-6 in the worm *C. elegans* and in human cell lines leads to centriole duplication failure and the consequent loss of centrioles, whereas in the algae *C. Reinhardtii* and flies *D. melanogaster*, mutations in *Sas-6* result in centriole symmetry aberrations (Leidel *et al*., 2005; Nakazawa *et al*, 2007; Rodrigues-Martins *et al*, 2007).

Intriguingly, in early mouse development, centrioles form *de novo* around the blastocyst stage (embryonic day (E) 3.5) (Courtois *et al*, 2012; Gueth-Hallonet *et al*, 1993), from which mouse embryonic stem cells (mESCs) are derived and propagated. To our knowledge, there are currently no reports on the roles of SAS-6 in centriole assembly or integrity in mice or mouse cells. In this study, we asked whether SAS-6 is required for centriole *de novo* formation and duplication during mouse development and in mESCs. Our data show that the loss of SAS-6 in the developing mouse leads to centriole formation failure, activation of the 53BP1-, USP28- and p53-dependent mitotic surveillance cell death pathway, and arrest of development at mid-gestation (E9.5). In contrast, mESCs without SAS-6 can still form centrioles, which are nonetheless structurally defective and lack the capacity to template cilia. While *Sas-6* null blastocyst cells acquire centrioles in culture, mESCs exit from pluripotency leads to the loss of centrioles. Our data indicate that mESCs possess a remarkable ability to bypass the requirement for SAS-6 in centriole duplication in other cellular contexts and highlights the importance of considering the species differences and cellular differentiation status even for house-keeping functions like centriole duplication, spindle organization and cell division.

## Results and Discussion

### Mutation in mouse *Sas-6* leads to embryonic arrest around mid-gestation

To determine the functions of mouse SAS-6 *in vivo*, we used CRISPR/Cas9 to generate *Sas-6* knockout mice by targeting exon 4 (Materials and Methods, Table 1). The resulting *Sas-6* mutant allele (*Sas-6*^*em4/em4*^) had a frameshifting deletion, which is predicted to lead to a premature stop codon (Table 1). *Sas-6*^*em4/em4*^ embryos arrested development ∼E9.5, when they still formed a heart but did not show somites or undergo embryonic turning that are typical in wild-type (WT) embryos (Fig.1A, Fig. S1A). The phenotype of *Sas-6*^*em4/em4*^ embryos resembled our previously reported *Sas-4*^*−/−*^ embryos without centrioles (Bazzi & Anderson, 2014a), suggesting a crucial role for SAS-6 in centriole formation and mouse development.

**Table 1.**
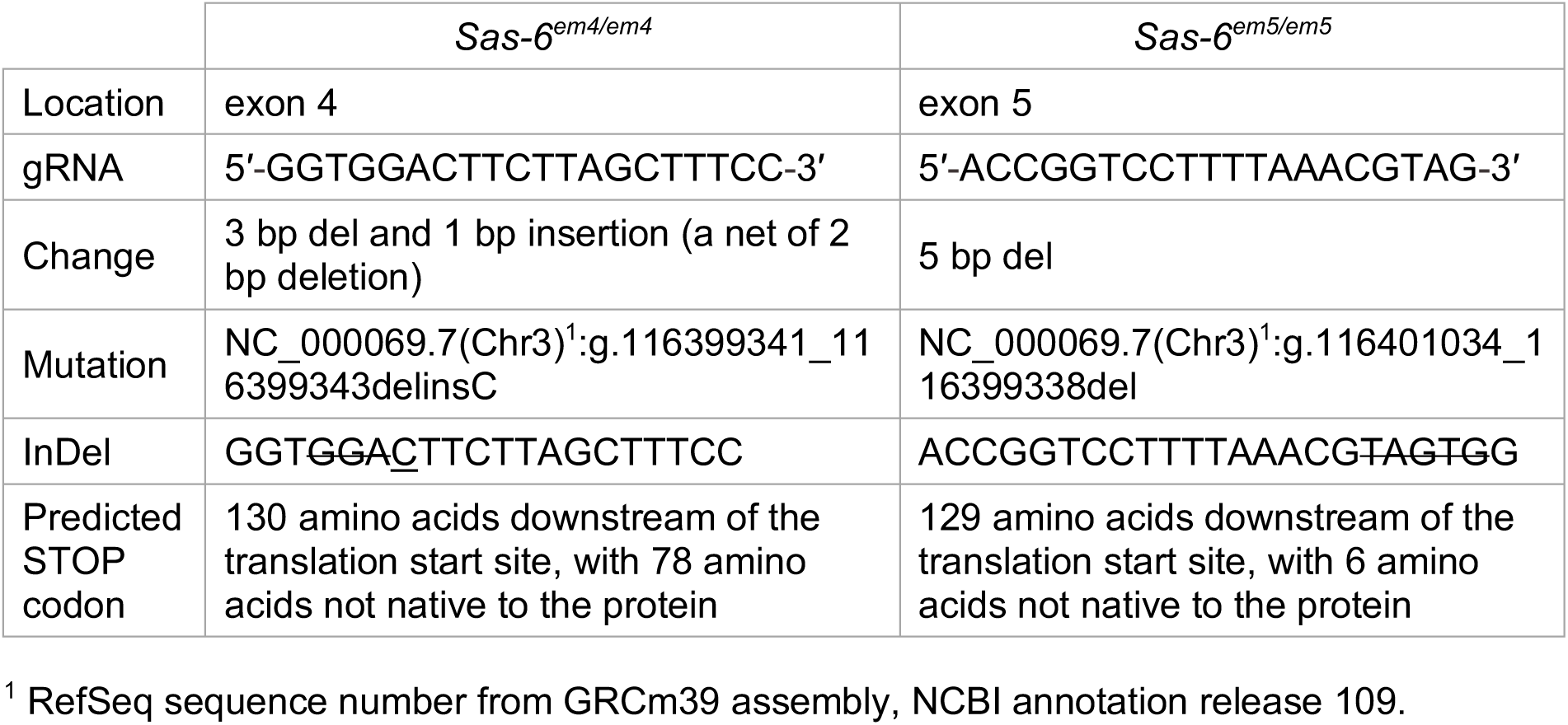
Description of CRISPR/Cas9 mediated knockouts of *Sas-6* in the mouse *in vivo*.

**FIG. 1.**
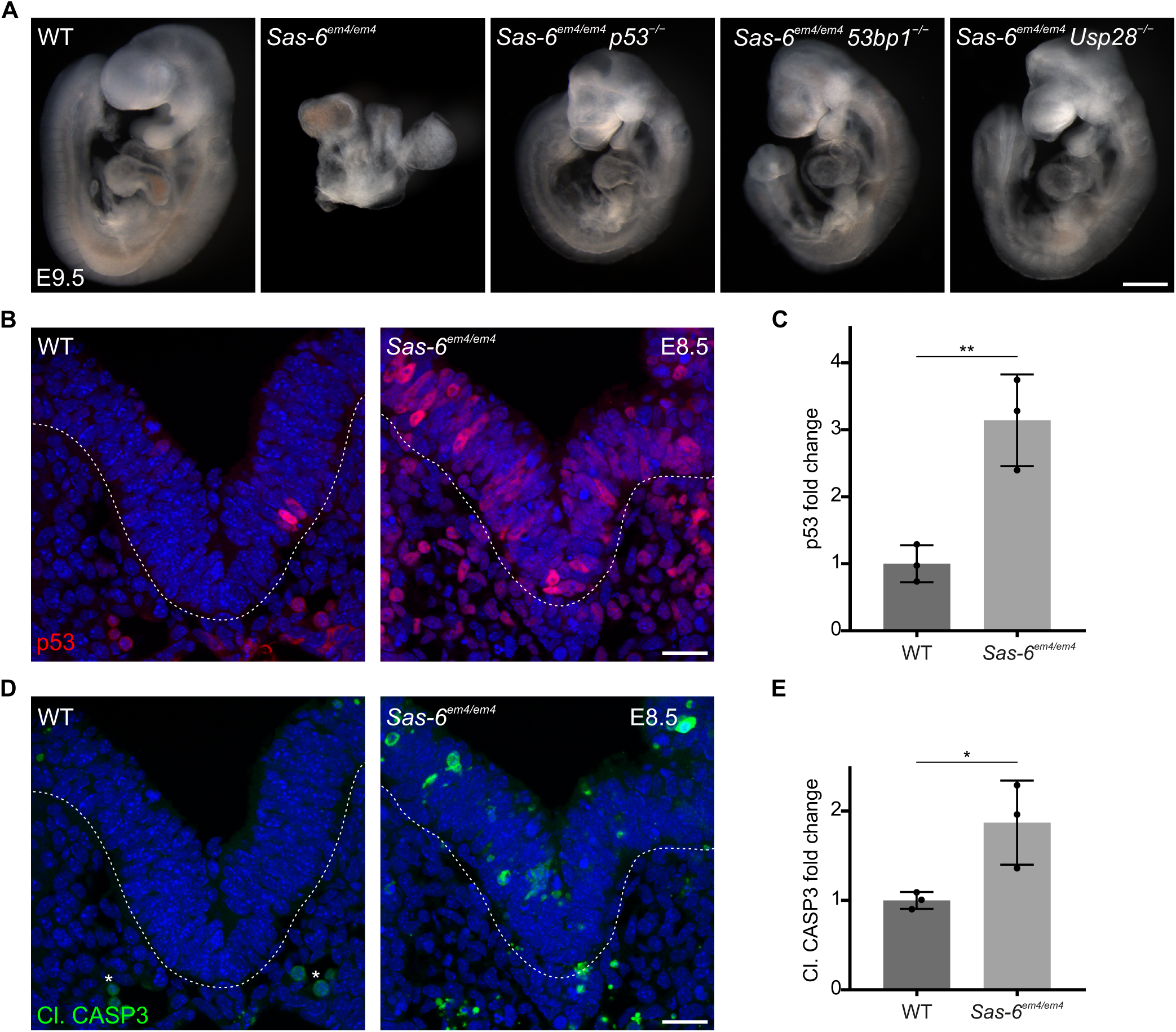
Mutation in mouse SAS-6 activates the 53BP1-USP28-p53 mitotic surveillance pathway. A. WT, *Sas-6*^*em4/em4*^, *Sas-6*^*em4/em4*^ *p53*^− /−^, *Sas-6*^*em4/em4*^ *53bp1*^− /−^ and *Sas-6*^*em4/em4*^ *Usp28*^− /−^ embryos at E9.5. At least three embryos per genotype showed similar phenotypes. Scale bar = 500 µm. B. Immunostaining for p53 on transverse sections of WT and *Sas-6*^*em4/em4*^ embryos at E8.5. The sections shown encompasses the neural plate (top) and mesenchyme (bottom), demarcated by the dashed line. Asterisks indicate non-specific staining of blood cells. Scale bars = 25 µm. C. Quantification of the nuclear p53 in (B). Three embryos per genotype were used for the quantifications. Values were normalized to WT. Error bars represent mean ± s.d. WT: 1.00 ± 0.23 (n = 2672 cells); *Sas-6*^*em4/em4*^: 3.14 ± 0.56 (n = 2758). **P < 0.01 (two-tailed Student’s t-test). D. Immunostaining for Cleaved-Caspase3 (Cl. CASP3) as mentioned in (B). E. Quantification of Cl. CASP3 in (D) as mentioned in (C). WT: 1.00 ± 0.08 (n = 2672); *Sas-6*^*em4/em4*^: 1.87 ± 0.38 (n = 2758). *P < 0.05.

### The loss of SAS-6 activates the 53BP1-USP28-p53 mitotic surveillance pathway

In order to assess whether the loss of SAS-6 leads to p53 upregulation and cell death, as in *Sas-4*^−/−^ mutants (Bazzi & Anderson, 2014a), we performed immunostaining for p53 and active cleaved-caspase 3 (Cl. CASP3) on sections from WT and *Sas-6*^*em4/em4*^ embryos at E8.5 (Fig. 1B, D). In comparison to WT embryos where both were rarely detectable, the *Sas-6*^*em4/em4*^ mutants showed significantly increased levels of nuclear p53 (∼3-fold) and Cl. CASP3 (∼2-fold) (Fig.1B-E).

Next, we asked whether the *Sas-6* mutant phenotype is caused by the activation of the p53-, 53BP1- and USP28-dependent mitotic surveillance pathway (Xiao *et al*., 2021). To functionally address this question, we crossed *Sas-6*^+/*em4*^ mice to *p53*^+/−^, *53bp1*^+/−^ or *Usp28*^+/−^ null mouse alleles (Marino *et al*, 2000; Xiao *et al*., 2021). All three double mutant embryos: *Sas-6*^*em4/em4*^ *p53*^−/−^, *Sas-6*^*em4/em4*^ *53bp1*^−/−^ and *Sas-6*^*em4/em4*^ *Usp28*^−/−^, were evidently rescued at E9.5 as judged by the normalized size and morphology, which were more similar to WT when compared to the *Sas-6*^*em4/em4*^ single-mutant embryos (Fig.1A). In this regard, the double mutant embryos showed body turning and somites, and were also similar to our reported mitotic surveillance pathway double mutants with *Sas-4* (Xiao *et al*., 2021). The data indicated that, similar to SAS-4, the loss of SAS-6 in the mouse activates the mitotic surveillance pathway leading to cell death and embryonic arrest at mid-gestation.

### Mouse SAS-6 is essential for centriole formation *in vivo*

To examine whether SAS-6 is required for centriole formation in the mouse, we stained embryo sections at E9.0 for the centrosomal marker γ-tubulin (TUBG) and a mother centriole distal appendage marker (CEP164). We detected centrioles, as defined by the co-localization of TUBG and CEP164, in almost all of the cells in WT embryos (97%). In contrast, centrioles were identified only in a minor fraction of cells in *Sas-6*^*em4/em4*^ mutants (16%) (Fig. S1B, C). Intriguingly, when we used the same criteria to analyze an earlier developmental stage ∼E7.5, we only rarely detected centrioles in *Sas-6*^*em4/em4*^ mutants at this stage (1%) (Fig. S1D, E).

These findings prompted us to reassess whether the residual centrosomes in *Sas-6*^*em4/em4*^ embryos formed after E7.5 due to the allele being hypomorphic for *Sas-6*. Thus, we generated another *Sas-6* mutant allele with a frameshifting deletion in exon 5 (*Sas-6*^*em5/em5*^), which is predicted *in silico* to result in a premature stop codon (Table 1). In contrast to *Sas-6*^*em4/em4*^, immunostaining of *Sas-6*^*em5/em5*^ embryos at E9 for TUBG and CEP164 did not detect any centrioles (Fig. S1B, C). Instead, in a minor fraction of interphase cells we observed focal accumulations of TUBG that did not co-localize with CEP164 signal (12%) (Fig. S1B, C), suggesting that these accumulations were most likely devoid of mother centrioles (see below). Immunostaining for SAS-6 and TUBG confirmed that SAS-6 co-localized with TUBG in almost all cells of WT embryo sections (95%) at E9.0, in 19% of cells in *Sas-6*^*em4/em4*^ and only rarely in *Sas-6*^*em5/em5*^ (4%) (Fig. S1F, G).

For higher resolution analyses of centriole formation in *Sas-6* mutant embryos, we utilized ultra-expansion microscopy (U-ExM), a technique that relies on isometrically expanding the sample ∼4 times and has been recently widely implemented for centriole analyses (Gambarotto *et al*, 2019). We combined U-ExM with immunostaining for the centriolar wall marker, acetylated tubulin (Ac-TUB). We observed that in WT embryo sections (E9.0), each pole of a mitotic spindle contained a pair of centrioles, while in *Sas-6*^*em4/em4*^ mutants, only rare and single centrioles were detected (11%), suggesting centriole duplication failure (Fig. S1H, I). Of note, no centrioles were detected in the mitotic poles of *Sas-6*^*em5/em5*^ embryos (Fig. S1H, I). Importantly, the phenotype of *Sas-6*^*em5/em5*^ at E9.5 embryos was almost identical to that of *Sas-6*^*em4/em4*^ (Fig. S1A). The data suggested that the *Sas-6*^*em4/em4*^ is a severe hypomorphic allele of *Sas-6*, whereas the *Sas-6*^*em5/em5*^ is a null allele of *Sas-6*, and that SAS-6 is essential for centriole formation in mouse embryos *in vivo*.

### SAS-6 is required for centriole integrity in mESCs

To study the roles of *Sas-6* in an *in vitro* setting that mimics mouse embryonic development, we chose to knockout *Sas-6* in mESCs. To accomplish this without any reasonable doubt of residual SAS-6 protein, we used CRISPR/Cas9 with a pair of guide RNAs (gRNAs) flanking the open reading frame (ORF) of *Sas-6*, and engineered a null allele lacking the entire *Sas-6* ORF (*Sas-6*^*−/−*^) (Fig. S2A, Materials and Methods, Table 2). The deletion of the *Sas-6* ORF was validated by PCR and Western blot (Fig. S2B, C). We next used immunostaining for TUBG to assess centrosome formation in *Sas-6*^*−/−*^ mESCs. While centrosomes were evident in nearly every cell in WT mESCs (93%), it was surprising that the majority of *Sas-6*^*−/−*^ mESCs also showed apparent centrosomes (87%) (Fig. 2A, B). This unexpected observation suggested that unlike in mouse embryos, *Sas-6* may not be essential for centrosome formation in mESCs.

**Table 2.**
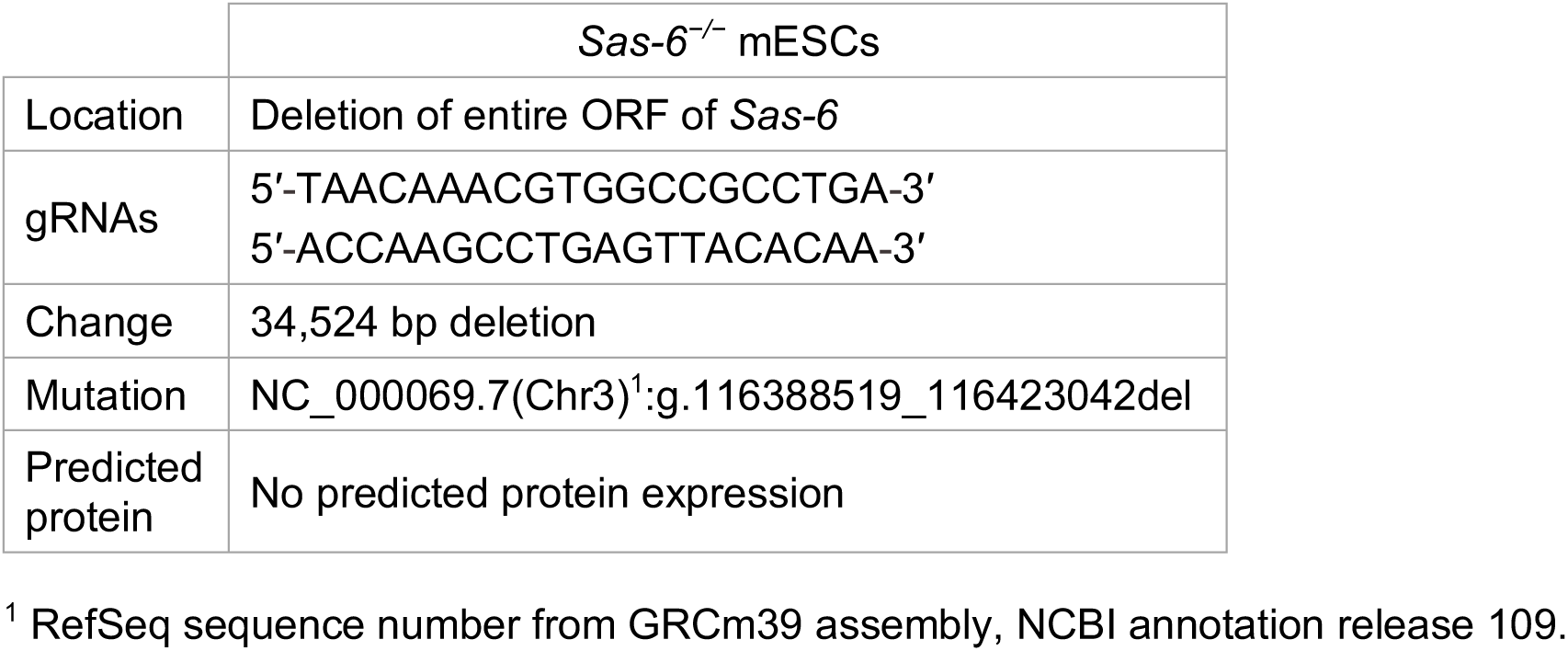
Description of CRISPR/Cas9 mediated knockout of *Sas-6* in mESCs.

**FIG. 2.**
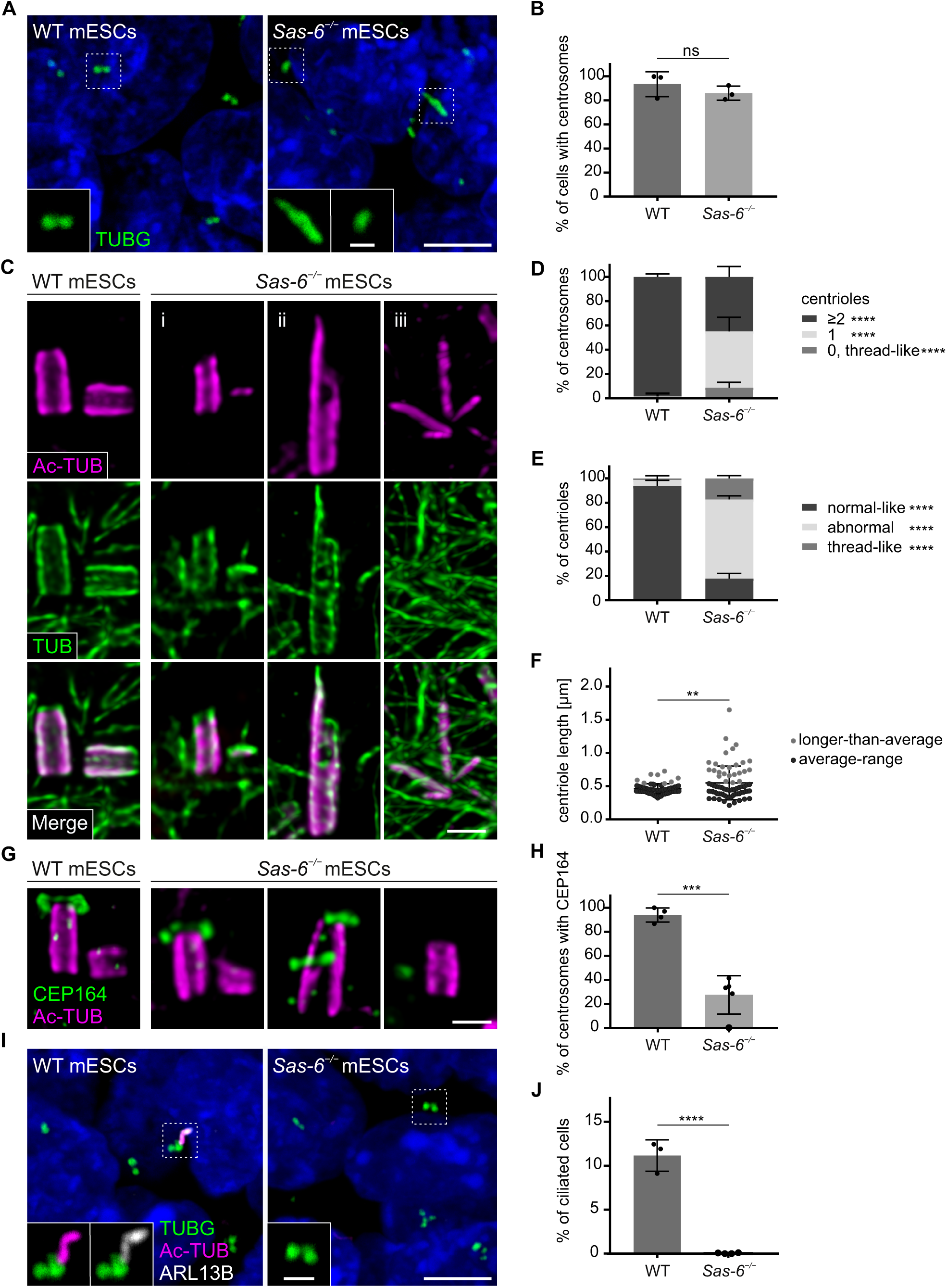
SAS-6 is required for centriole integrity and cilia formation in mESCs. A. Immunostaining for TUBG in WT and *Sas-6*^*−/−*^ mESCs. Insets are magnifications of the dashed squares. Scale bars = 5 µm and 1 µm (insets). B. Quantification of the percentage of cells with centrosomes (TUBG) in (A) from three independent experiments. Error bars represent mean ± s.d. WT: 93 ± 8% (n = 1679 cells); *Sas-6*^*−/−*^ : 87 ± 5% (n = 1168). ns = not significant with p > 0.05 (two-tailed Student’s t-test). C. Centrioles visualized using U-ExM and immunostaining for α- and β-tubulin (TUB) and Ac-TUB in WT and *Sas-6*^*−/−*^ mESCs: (i) normal-like centrioles, (ii) abnormal centrioles, (iii) thread-like structures. Scale bars = 300 nm. D. Quantification of the percentage of centrosomes with ≥ 2, 1, or 0 centrioles in (C) in WT and *Sas-6*^*−/−*^ mESCs from five independent experiments. Error bars represent mean ± s.d. WT (n = 156 centrosomes): ≥ 2 centrioles = 99 ± 2%; 1 centriole = 1 ± 2%; *Sas-6*^*−/−*^ (n = 254): ≥2 centrioles = 45 ± 8%, 1 centriole = 46 ± 10%, 0 centrioles = 9 ± 4%. ***P < 0.001, *P < 0.05 (two-tailed Student’s t-test) E. Quantifications of the percentage of centrioles within each category in (C) from five independent experiments. Error bars represent mean ± s.d. WT (n = 330 centrioles): normal-like centrioles = 94 ± 4%; abnormal centrioles = 5 ± 3%; thread-like structures = 1 ± 2%; *Sas-6*^*−/−*^ (n = 432): normal-like centrioles = 18 ± 4%, abnormal centrioles = 65 ± 3%, thread-like structures = 17 ± 2%. ****P < 0.0001, (two-tailed Student’s t-test). F. Quantification of centriole length of normal-like centrioles in (C) in WT and *Sas-6*^*−/−*^ mESCs from five independent experiments. Data points higher than the mean + 1 s.d of WT are defined as longer-than-average centrioles (gray dots). Error bars represent mean ± s.d. WT: 0.46 ± 0.07 µm (n = 72 centrioles); *Sas-6*^*−/−*^ : 0.55 ± 0.25 µm (n = 72). ***P < 0.001, *P < 0.05 (two-tailed Student’s t-test). G. Immunostaining for Ac-TUB and CEP164 in U-ExM of WT and *Sas-6*^*−/−*^ mESCs. Scale bars = 300 nm. H. Quantification of the percentage of centrosomes with mother centrioles (Ac-TUB) with the distal appendage marker (CEP164) in (G). Error bars represent mean ± s.d. WT: 94 ± 5% (n = 104 centrosomes from four independent experiments); *Sas-6*^*−/−*^ : 28 ± 14% (n = 140 from five experiments). ***P < 0.001 (two-tailed Student’s t-test). I. Immunostaining of the cilia markers ARL13B and Ac-TUB, while basal bodies are marked with TUBG, on WT and *Sas-6*^*−/−*^ mESCs. The insets show separate channels for selected basal bodies and cilia magnified from the dashed squares. Scale bars = 5 µm and 1 µm (insets). J. Quantification of the percentage of ciliated cells in (I). Error bars represent mean ± s.d. WT: 11 ± 1% (n = 2602 cells from three experiments); *Sas-6*^*−/−*^ : 0 ± 0% (n = 4602 from four experiments). ****P < 0.0001 (two-tailed Student’s t-test).

To assess whether the TUBG-marked centrosomes in *Sas-6*^*−/−*^ mESCs contained centrioles at their core, we used U-ExM combined with immunostaining for *bona fide* centriolar markers (Ac-Tub and α/β-tubulin, TUB) (Fig. 2C). Almost all of the centrosomes in WT cells contained two or more centrioles (99%) and only a rare fraction contained one centriole (1%) (Fig. 2D). In contrast, in *Sas-6*^*−/−*^ mESCs, almost half of the centrosomes had two or more centrioles (46%), and a comparable fraction had one centriole (45%) (Fig. 2D). Additionally, in a minor fraction of *Sas-6*^*−/−*^ centrosomes, aberrant centriolar threads were detected (9%) (Fig 2C, D). We classified the centriolar structures in *Sas-6*^*−/−*^ mESCs into the following three categories: i) normal-like centrioles (18%), ii) abnormal centrioles (65%), and iii) thread-like structures (17%) (Fig. 2C, E). To gain a more quantitative insight into the structure of the centrioles in *Sas-6*^*−/−*^ mESCs, we measured the length of the normal-like centrioles and found that they were significantly longer in in *Sas-6*^*−/−*^ than in WT (Fig. 2F). In this respect, around one third of these centrioles were elongated (35%), compared to a minor fraction of longer-than-average centrioles in WT mESCs (13%) (gray dots in Fig. 2F). In summary, the data suggested that *Sas-6*^*−/−*^ centrioles in mESCs had a compromised ability to duplicate and/or were unstable with mostly abnormal structures.

### The centrioles in *Sas-6*^*−/−*^ mESCs have proximal and distal defects

SAS-6 has been shown to cooperate with STIL, an essential component for centriole duplication, to initiate procentriole formation (Kratz *et al*, 2015). To address whether the fraction of centrioles that failed to duplicate in *Sas-6*^*−/−*^ mESCs is associated with an impairment of STIL recruitment, we used U-ExM combined with TUB, Ac-TUB and STIL immunostaining (Fig. S2D). STIL localized to the majority of centrosomes in WT cells (74%), which is consistent with percentage of cells undergoing centriole duplication during the S/G2/M phases of the cell cycle in mESCs (∼70% in asynchronous cells) (Fig. S2D, E). In *Sas-6*^*−/−*^ mESCs, STIL localized to less than one third of the centrosomes (29%) (Fig. S2D, E), which is roughly half of the STIL-positive centrosomes in WT cells, and might account for the duplication failure in almost half of the *Sas-6*^*−/−*^ centrosomes (single centrioles in Fig. 2D).

To assess whether the loss of centriole integrity in *Sas-6*^*−/−*^ mESCs is associated with disruption of the proximal centriole end or internal structural scaffold, we immunostained centrioles in U-ExM for CEP135 (proximal end) and POC5 (scaffold). We found that CEP135 localized to the proximal centriole in all WT cells, while the number of centrioles with CEP135 was slightly decreased in *Sas-6*^*−/−*^ cells (73%) (Fig. S2F, G). Notably, only a minority of CEP135-positive centrioles in *Sas-6*^*−/−*^ mESCs showed normal CEP135 localization (12%) (Fig. S2F, H). On the other hand, POC5 was present in WT and *Sas-6*^*−/−*^ intact centrioles, but also lining the abnormal centrioles and centriolar threads in *Sas-6*^*−/−*^ cells, further confirming their centriolar nature (Fig. S2I).

To examine whether the mother centrioles in *Sas-6*^*−/−*^ mESCs were decorated with distal appendages, we used U-ExM combined with Ac-TUB and CEP164 immunostaining. The data showed, as expected, that CEP164 mostly localized to the mother centrioles in WT centrosomes (94%) (Fig. 2G, H). In contrast, only a quarter of the centrosomes in *Sas-6*^*−/−*^ mESCs had centrioles associated with CEP164 (28%) (Fig. 2G, H). To determine whether the abnormal centrioles in *Sas-6*^*−/−*^ mESCs retained the ability to template cilia, we used immunostaining with the ciliary axoneme marker Ac-TUB and ciliary membrane protein ARL13B. Although cilia were present in only a small fraction of WT mESCs (11%), no cilia were detected in *Sas-6*^*−/−*^ mESCs (Fig. 2I, J), suggesting that SAS-6 is not only required for centriole integrity, but also distal appendage recruitment and cilia formation in mESCs.

### Short-term culture of *Sas-6*^*em5/em5*^ null blastocysts induces centriole formation

Because mESCs are usually derived from mouse blastocysts (E3.5), we asked whether SAS-6 is essential for the *de novo* formation of centrioles, which starts ∼E3. To unambiguously identify centrioles *in vivo*, we crossed the *Sas-6*^*+/em5*^ null allele to *Cetn2-eGFP*^*+/tg*^, a transgenic mouse line with centrioles marked with centrin-eGFP (Bangs *et al*, 2015; Higginbotham *et al*, 2004), and examined *Sas-6*^*em5/em5*^ *Cetn2-eGFP*^*+/tg*^ blastocysts immunostained for centrosomes using TUBG (Fig. 3A). The majority of the cells in control *Cetn2-eGFP*^*+/tg*^ blastocysts had foci positive for both markers (73%) (Fig. 3A, B). In contrast, in mutant *Sas-6*^*em5/em5*^ *Cetn2-eGFP*^*+/tg*^ blastocysts, only minor TUBG accumulations were observed (23%), but they did not contain CETN2-eGFP, suggesting that they are devoid of centrioles (Fig. 3A, B). The data confirmed that *Sas-6*^*em5/em5*^ null mutants do not seem to form centrioles in early development.

**FIG. 3.**
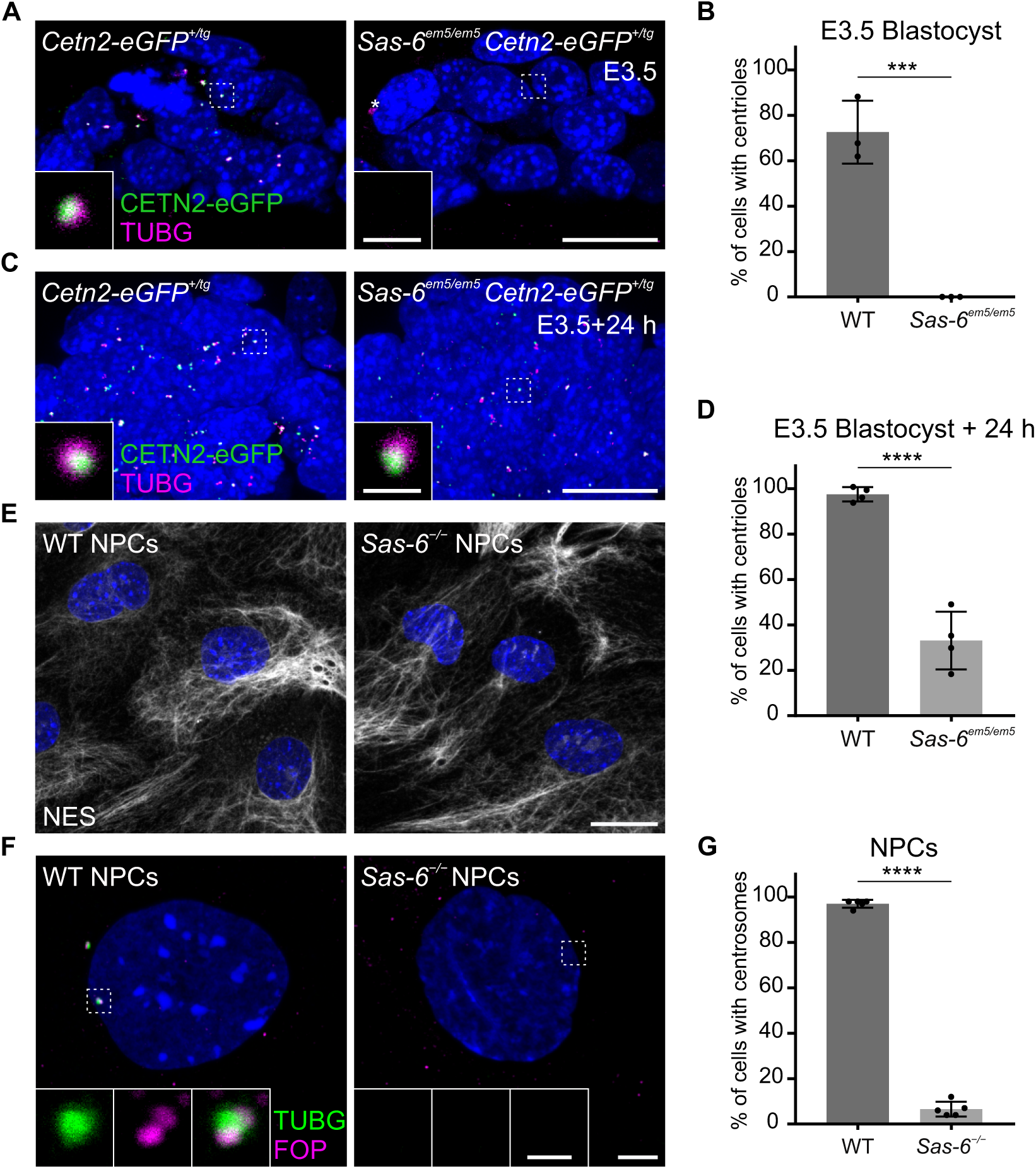
Centrioles in *Sas-6*^*−/−*^ mESCs are formed *de novo* during the derivation from blastocysts and are lost during differentiation. A. Whole-mount immunostaining for TUBG on *Cetn2-eGFP*^*+/tg*^ and *Sas-6*^*em5/em5*^ *Cetn2-eGFP*^*+/tg*^ blastocysts at E3.5. Insets are magnifications of the dashed squares. Scale bars = 20 µm and 1 µm (insets). B. Quantification of the percentage of cells with centrioles (Centrin-eGFP) in E3.5 blastocysts in (A). Three blastocysts per genotype were used for the quantifications. Error bars represent mean ± s.d. (C) WT: 73 ± 11% (n = 200 cells); *Sas-6*^*em5/em5*^: 0 ± 0% (n = 175). ***P < 0.001, (two-tailed Student’s t-test). C. Whole-mount immunostaining as mentioned in (A) on blastocysts after 24 h of culture. D. Quantification as mentioned in (B) from blastocysts after 24 h of culture in (C). Four blastocysts per genotype were used for the quantifications. WT: 98 ± 30% (n = 630 cells); *Sas-6*^*em5/em5*^: 33 ± 11% (n = 690). ****P < 0.0001. E. Immunostaining for Nestin (NES) on WT and *Sas-6*^*−/−*^ cells after *in vitro* neural differentiation of mESCs. Scale bar = 20 µm. F. Immunostaining for TUBG and FOP in WT and *Sas-6*^*−/−*^ cells after *in vitro* neural differentiation. The insets show the separate channels for the magnified areas in the dashed squares. Scale bars = 5 µm and 1 µm (insets). G. Quantification of the percentage of cells with centrosomes (TUBG and FOP) in (F) from five independent experiments. Error bars represent mean ± s.d. WT: 97 ± 0% (n = 1388 cells); *Sas-6*^*−/−*^ : 6 ± 0% (n = 1068). ****P < 0.0001, (two-tailed Student’s t-test).

Given that *Sas-6*^*em5/em5*^ null embryos at E3.5 do not form centrioles but CRISPR-generated *Sas-6*^*−/−*^ mESCs still manage to form abnormal centrioles, we next asked whether centrioles can form *de novo* during the derivation of mESCs from the *Sas-6*^*em5/em5*^ *Cetn2-eGFP*^*+/tg*^ acentriolar blastocysts. After 24 hours (h) of culture, almost all of the cells in the cultured WT blastocysts contained centrioles (98% with both CETN2-eGFP and TUBG), and surprisingly, centrioles were already detectable in one third of the cells in *Sas-6*^*em5/em5*^ *Cetn2-eGFP*^*+/tg*^ blastocysts (33%) (Fig. 3C, D). The data suggested that the mESC culture conditions are conducive to *de novo* centriole formation in the absence of SAS-6.

### The differentiation of *Sas-6* mutant mESCs leads to centriole loss

To test whether the ability to form centrioles, albeit mostly abnormal, via a SAS-6-independent pathway is a characteristic of the pluripotent mESCs, we analyzed centriole formation in mESCs differentiated and enriched for neural progenitor cells (NPCs). As expected, centrosomes immunostained for TUBG and FOP (a centriole marker) were detected in almost all WT NPCs expressing the intermediate filament NESTIN (NES) (97%, Fig. 3E-G). Remarkably, in *Sas-6*^*−/−*^ NPCs, the number of cells with centrosomes sharply decreased upon differentiation (from 87%, Fig. 2B, down to 6%, Fig. 3G). The data suggested that the SAS-6-independent centriole formation pathway is a property of pluripotent mESCs that is largely lost upon differentiation.

In this study, we report that mutations in mouse *Sas-6* cause embryonic arrest at mid-gestation with elevated levels of p53 and cell death, as well as the activation of the p53-, 53BP1- and USP28-dependent mitotic surveillance pathway. We have previously reported similar phenotypes of elevated p53 and cell death for other genes essential for centriole duplication, such as *Sas-4* and *Cep152* (Bazzi & Anderson, 2014a). The current data demonstrated that mouse SAS-6 is required for centriole formation in developing mouse embryos (Fig. 4), as expected from the established role of its orthologs in *C. elegans* and human cells (Gupta *et al*, 2020; Leidel *et al*., 2005; Wang *et al*, 2015). Together, the data provide further support that the activation of the mitotic surveillance pathway is not specific to the loss of specific centriolar proteins but the centriole/centrosome structure *per se*.

**FIG. 4.**
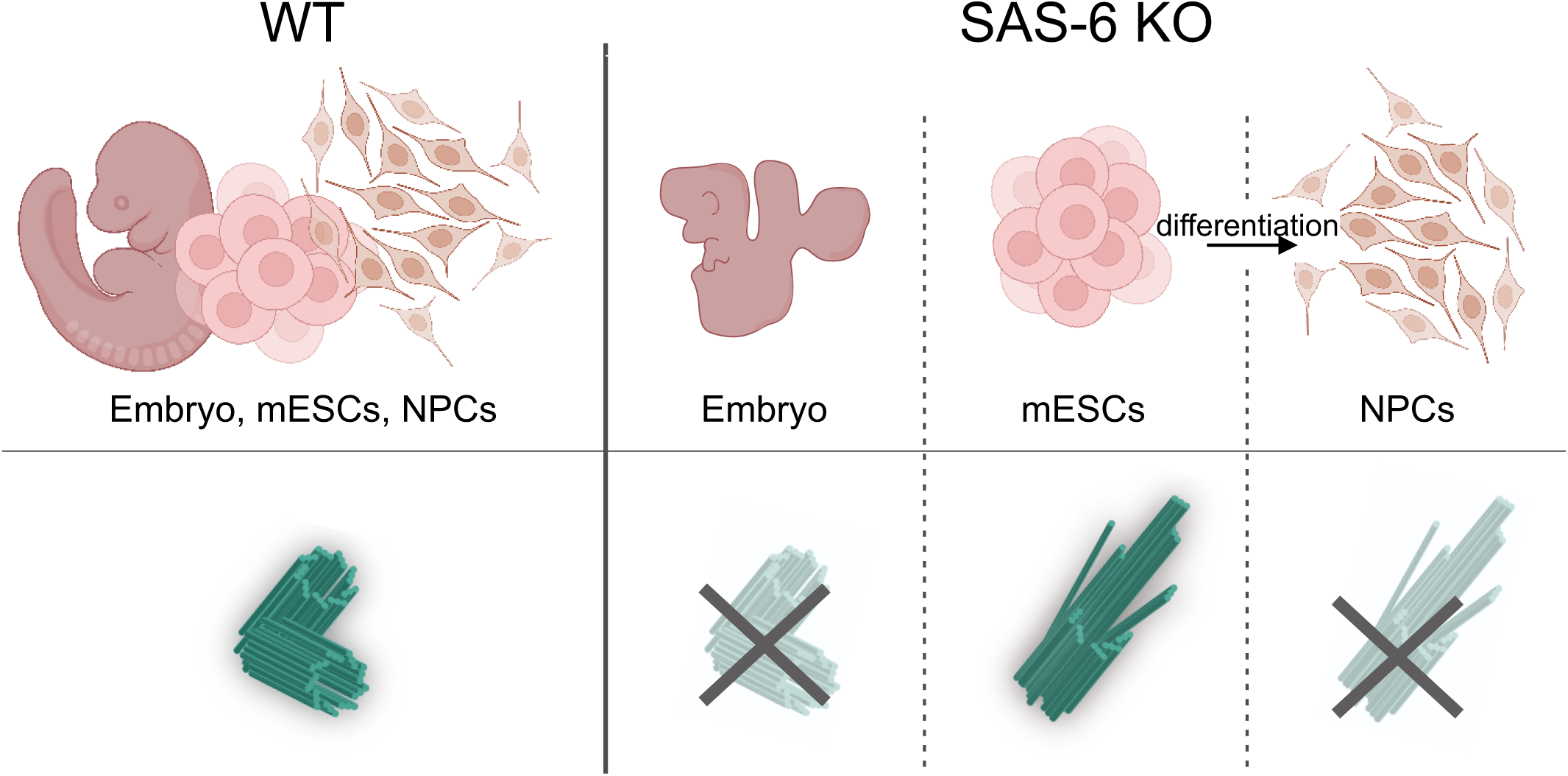
Graphical summary depicting the roles of SAS-6 in mouse embryos, mESCs and NPCs. The loss of SAS-6 in the developing mouse leads to centriole formation failure. In contrast, SAS-6 is not strictly required for centriole formation in mESCs, but is still important to regulate centriole structure and integrity. *Sas-6* mutant mESC exit from pluripotency and differentiation into neural progenitors (NPCs) leads to the loss of centrioles. Figure was created with BioRender.com.

To our surprise, and in contrast to the mouse embryo, removing SAS-6 from mESCs is still compatible with the formation of centrioles, but these centrioles are mostly abnormal and defective (Fig. 4). Our data support published literature on SAS-6-independent centriole duplication that also leads to the formation of abnormal centrioles in evolutionarily more distant organisms, namely *C. Reinhardtii* and *D. melanogaster* (Nakazawa *et al*., 2007; Rodrigues-Martins *et al*., 2007). Interestingly, we demonstrate that centrioles appear during the derivation process in SAS-6 deficient mESCs but are lost again upon differentiation (Fig. 4). The findings extend the observations on abnormal centrioles upon SAS-6 depletion and reveal the differential requirements for SAS-6 even within cells of the same species. Our work sheds light on cell type specific functions and composition of centrosomes, particularly in stem cells (Fig. 4).

In *D. melanogaster, C. elegans* and human cells, SAS-6 has been shown to directly interact with a downstream protein STIL, and provide a structural basis for the recruitment of other centriole duplication proteins, such as CEP135 and SAS-4 (Lin *et al*, 2013; Qiao *et al*, 2012; Tang *et al*, 2011). The loss of STIL or SAS-4 leads to centriole duplication failure in all organisms and cell types studied to date (Bazzi & Anderson, 2014a; Vulprecht *et al*, 2012). Notably, unlike SAS-6, SAS-4 is essential for centriole formation in mESCs (Xiao *et al*., 2021). Our data show that STIL localized to half of the centrosomes in *Sas-6*-null mESCs, which might account for centriole duplication failure in almost half of the centrosomes that contain only single centrioles. Our data suggest that pluripotent mESCs without SAS-6 have a bypass pathway of SAS-4 recruitment, which may be compensated for by another factor or an unidentified SAS-6 homolog.

Although centrioles are still present in *Sas-6*^*−/−*^ mESCs, they exhibit profound proximal and distal defects. Whether this phenotype arises as a consequence of an improper initiation of assembly or a later destabilization of the microtubule triplets, are still open questions. The cartwheel protein, CEP135, has been shown to be important for centriole stability in *T. thermophila* and human cells (Bayless *et al*, 2012; Lin *et al*., 2013). Our data show that centrioles in *Sas-6*^*−/−*^ mESCs exhibit abnormal localization of CEP135. We speculate that the lack of an initial stable cartwheel scaffold that provides a 9-fold symmetry template and the ensuing mis-localization of CEP135 may account for the abnormal centriole architecture and its instability.

In conclusion, our work provides new fundamental insights into alternative and SAS-6-independent pathways of centriole formation in mammalian cells and shows that mESCs are a special *in vitro* model for centriole biology that is akin to the more “primitive” origins of life such as algae.

## Materials and Methods

### Mice and genotyping

The following mouse alleles for *Usp28*^*+/-*^ (*Usp28*^*em1/Baz*^) and *53bp1*^*+/-*^ (*Trp53bp1*^*em1/Baz*^) (Xiao, 2021), as well as *Cetn2-eGFP*^*+/tg*^ (Tg^(CAG-EGFP/CETN2)3-4Jgg^) (Bangs *et al*., 2015; Higginbotham *et al*., 2004) were used in this study. The *p53*^*+/-*^ (*Trp53*^*+/-*^) null allele was generated by crossing *Trp53*^*+/tm1*.*1Brn*^ (Jax stock no. 008462, Marino, 2000) mice with *K14-Cre*^*+/tg*^ (Hafner *et al*, 2004) females, which express the Cre recombinase in the zygote. *Sas-6*^*+/em4*^ (*Sass6*^*em4/Baz*^, in exon 4) and *Sas-6*^*+/em5*^ (*Sass6*^*em5/Baz*^, in exon 5) mice were generated using CRISPR/Cas9 genome-editing by the CECAD *in vivo* Research Facility (ivRF, Branko Zevnik) (Table 1), where gRNA, Cas9 protein and mRNA were delivered to fertilized zygotes by pronuclear injection or electroporation (Chu *et al*, 2016; Troder *et al*, 2018). The animals were housed and bred under SOPF conditions in the CECAD animal facility. The animal generation (84-02.04.2014.A372), notifications and breeding applications (84-02.05.50.15.039, 84-02.04.2015.A405, UniKöln_Anzeige§4.20.026, 84-02.04.2018.A401, 81-02.04.2021.A130) were approved by the Landesamt für Natur, Umwelt, und Verbraucherschutz Nordrhein-Westfalen (LANUV-NRW) in Germany. The phenotypes were analyzed in the FVB/NRj background. Genotyping was carried out using standard and published PCR protocols as cited or described in this work. For the new *Sas-6*^*+/em4*^ and *Sas-6*^*+/em5*^ mouse alleles, the PCR products (primers shown in Table S1) were digested with *AvaII* and *HpyCH4IV* restriction enzymes (New England BioLabs; Ipswich, MA, USA), respectively, to distinguish between the WT and mutant alleles. *AvaII* cut the product in the *Sas-6*^*+/em4*^ mutant allele, whereas *HpyCH4IV* did not cut the product in the *Sas-6*^*+/em5*^ mutant allele.

### Mouse embryonic stem cell culture

mESCs were derived from WT blastocysts as previously described (Bryja *et al*, 2006). The established mESC lines were adapted to feeder-free conditions and cultured on 0.1% gelatin (PAN Biotech, Aidenbach, Germany) coated plates in Knock-Out DMEM (Thermo Fisher Scientific; Waltham, MA, USA) supplemented with 15% HyClone fetal bovine serum (FBS; VWR; Radnor, PA, USA), 2 mM L-glutamine (Biochrom; Berlin, Germany), 1% penicillin/streptomycin (Biochrom), 0.1 mM MEM non-essential amino acids (Thermo Fisher Scientific), 1 mM sodium pyruvate (Thermo Fisher Scientific), 0.1 mM β-mercaptoethanol (Thermo Fisher Scientific), 1000 U/ml leukemia inhibitory factor (LIF; Merck; Darmstadt, Germany), and with 1 μM PD0325901 (Miltenyi Biotec; Bergisch Gladbach, Germany) and 3 μM CHIR99021 (Miltenyi Biotec). Cells were maintained at 37°C with 6% CO2.

### Generation of *Sas-6* mutant mESCs using CRISPR/Cas9

For the generation of the CRISPR/Cas9-mediated *Sas-6* knockout mESCs line, a pair of gRNAs targeting the 5′ and 3′ ends of the entire *Sas-6* ORF (Fig. S2 and Table 2) were cloned as double-stranded oligo DNA into BbsI and SapI sites in pX330-U6-Chimeric_BB-CBh-hSpCas9 vector (Addgene; Watertown, MA, USA) modified with a Puro-T2K-GFP cassette containing puromycin-resistance by Dr. Leo Kurian’s research group (Center for Molecular Medicine Cologne). mESCs in suspension were transfected with the modified pX330 vector containing the pair of gRNAs using Lipofectamine 3000 (Thermo Fisher Scientific). The cells were then subjected to selection using puromycin (2 μg/ml, Sigma-Aldrich; St. Louis, MO, USA) 24h post-transfection for 2 days. After recovery for 5 days, individual colonies were picked and screened for the *Sas-6* locus deletion by PCR (Fig. S2 and Table 2). The cells were used for experiments after 4 passages.

### Embryo dissection, immunofluorescence staining and image acquisition

Timed pregnant female mice were sacrificed by cervical dislocation. Post-implantation embryos (E7.5-E9.5) were dissected in ice cold PBS with 0.1% Tween20 (AppliChem; Darmstadt, Germany), and fixed overnight in 2% paraformaldehyde (PFA; Carl Roth; Karlsruhe, Germany) at 4°C, or in methanol for 30 min at -20°C for Ultra-Expansion Microscopy (U-ExM) and SAS-6 immunostaining. Embryos were dehydrated overnight at 4°C in 30% sucrose and embedded in O.C.T. (Sakura Finetek; Alphen an den Rijn, Netherlands) for cryo-sectioning at 8 µm sections using a CM1850 Cryostat (Leica Biosystems; Wetzlar, Germany).

Pre-implantation E3.5 blastocysts were recovered by flushing the uterine horns with EmbryoMax M2 Medium (Sigma-Aldrich), and fixed in 4% PFA for 20 min at room temperature (RT) and 20 min at 4°C. Blastocysts were then permeabilized for 3 min with 0.5% Triton X-100 (Sigma-Aldrich) in PBS and three times for 10 min with immunofluorescence (IF) buffer containing 0.2% Triton X-100 in PBS. After blocking with 10% heat-inactivated goat serum in IF buffer, the blastocysts were incubated overnight with the primary antibodies at 4°C, followed by a 2h incubation with the secondary antibodies and DAPI at RT (1:1000, AppliChem). Blastocysts were imaged in single drops of PBS covered with mineral oil, followed by genotyping.

For IF staining of embryo sections, the slides were post-fixed in methanol for 10 min at -20°C, washed with IF buffer and blocked with 5% heat-inactivated goat serum in IF buffer. The slides were incubated overnight with primary antibodies at 4°C followed by 1h incubation with secondary antibodies and DAPI at RT (1:1000, AppliChem), then mounted with ProLong Gold Antifade reagent (Cell Signaling Technology; Danvers, MA, USA).

For IF staining of mESCs, the cells were cultured in Lab-Tek II chamber slides coated with 0.1% gelatin, fixed with 4% PFA for 10 min at RT and post-fixed with methanol for 10 min at -20°C. The cells were then permeabilized for 5 min using 0.5% Triton X-100 in PBS. After blocking with IF buffer with 5% heat-inactivated goat serum, the cells were incubated overnight with the primary antibodies at 4°C, followed by 1h incubation with secondary antibodies and DAPI (1:1000, AppliChem). The slides were mounted with ProLong™ Gold (Cell Signaling Technology).

Images were obtained using TCS SP8 (Leica Microsystems) and Stellaris 5 (Leica Microsystems) confocal microscopes.

### Antibodies

All primary antibodies are listed in Table 3. The following secondary antibodies were used: Alexa Fluor 488, 568, or 647 conjugates (Life Technologies) (IF 1:1000, U-ExM 1:400).

**Table 3.** List of primary antibodies used in this study.

| Antigen | Company | Catalog number | Dilution |
| --- | --- | --- | --- |
| p53 | Cell Signaling | 2524 | IF (1:2000) |
| Cl. CASP3 | Cell Signaling | 9661 | IF (1:400) |
| TUBG | Sigma-Aldrich | T6557 | IF (1:1000) |
| CEP164 | Proteintech | 22227-1-AP | IF (1:1000)<br>U-ExM (1:500) |
| SAS-6 | A kind gift from Renata Basto, Institut Curie, Paris, France |  | IF (1:300)<br>WB (1:1000) |
| Ac-TUB | Sigma-Aldrich | T6793 | IF (1:1000)<br>U-ExM (1:500) |
| α-TUB | Sigma-Aldrich | T6074 | U-ExM (1:500) |
| β-TUB | Sigma-Aldrich | T8328 | U-ExM (1:200 ) |
| ARL13B | Proteintech | 17711-1-AP | IF (1:1000) |
| STIL | Bethyl Laboratories | A302-441A | U-ExM (1:200) |
| CEP135 | Proteintech | 24428-1-AP | U-ExM (1:200) |
| POC5 | Bethyl Laboratories | A303-341A | U-ExM (1:250) |
| NES | BioLegend | PRB-315C | IF (1:500) |
| FOP | Sigma-Aldrich | WH0011116M1 | IF (1:500) |

### Ultra-Expansion Microscopy (U-ExM)

For U-ExM of mESCs, the cells were cultured on 12mm glass cover slips (VWR) coated with 0.1% gelatin and fixed with methanol for 10 min at -20°C. For U-ExM of embryos, cryo-sections were collected on 12mm glass coverslips, O.C.T. was washed away in PBS. Sample expansion was performed as described in (Gambarotto *et al*., 2019). Briefly, the cover slips were incubated in 1.4% formaldehyde (Sigma-Aldrich), 2% acrylamide (Sigma-Aldrich) in PBS for 5h at 37°C. Gelation was carried out in 35 µl of monomer solution (23% (w/v) sodium acrylate (Sigma-Aldrich), 10% (w/v) acrylamide, 0.1% (w/v) N,N′-methylenebisacrylamid (Sigma-Aldrich) in 1× PBS supplemented with 0.5% APS (Bio-Rad; Feldkirchen, Germany) and 0.5% TEMED (Bio-Rad) for 5 min on ice and 1h at 37°C, followed by gel incubation in denaturation buffer (200 mM sodium dodecyl sulfate (SDS; AppliChem), 200 mM NaCl and 50 mM Tris in H2O (pH 9) for 1.5h at 95°C, and initial overnight expansion in ddH_2_O at RT. The gels were incubated with primary antibodies for 3h at 37°C on an orbital shaker, then with secondary antibodies for 2.5h at 37°C. After the final overnight expansion in ddH_2_O at RT, the expanded gel size was accurately measured using a caliper, and then mounted on 12 mm glass cover slip coated with Poly-D-Lysine (Thermo Fisher Scientific) and imaged using a TCS SP8 confocal microscope with a Lightning suite (Leica Microsystems) to generate deconvolved images.

### Blastocysts *in vitro* culture

The blastocysts were cultured for 24h at 37°C and 6% CO_2_, in ibiTreat µ-Slides (Ibidi GmbH, Munich, Germany) on feeder cells that were previously growth-arrested with a 2h mitomycin C treatment (10 μg/ml, Sigma-Aldrich), in mESCs derivation media (mESCs media except with the FBS replaced with Knockout™ Serum Replacement (15%, Thermo Fisher Scientific). The cultures were fixed and processed for IF staining as described above for the E3.5 blastocysts. Slides were mounted with VectaShield mounting medium (Linaris; Dossenheim, Germany).

### Neural differentiation of mESCs

For neural differentiation, embryoid bodies were generated using the hanging drop method (Wang & Yang, 2008). Around 1000 mESCs were suspended in 20 µl of differentiation medium (mESCs media without LIF, PD0325901 and CHIR99021, supplemented with 1µM retinoic acid (Sigma-Aldrich). After 3 days, the resulting embryoid bodies were plated on ibiTreat µ-slides coated with fibronectin (30 μg/ml, Sigma-Aldrich) and cultured for 4 days. The cells were fixed in methanol for 10 min at -20°C for centrosomal staining or 4% PFA for 10 min at RT for other stainings, and processed for IF similar to mESCs. Slides were mounted with VectaShield mounting medium (Linaris).

### Western blotting

Western blots were performed according to standard procedures (Mahmood & Yang, 2012). Briefly, the cells or embryos were lysed in RIPA buffer (150 mM NaCl, 50 mM Tris pH 7.6, 1% Triton X-100, 0.25% sodium deoxycholate and 0.1% SDS (AppliChem) with an ethylenediaminetetraacetic acid (EDTA)-free protease inhibitor cocktail (Merck), phosphatase inhibitor cocktail sets II and IV (Merck), and phenylmethylsulfonyl fluoride (PMSF; Sigma-Aldrich). 80 µg of protein per sample was used for SDS-PAGE. Samples were then blotted onto polyvinylidene difluoride membrane (Merck). After blocking in 5% non-fat milk (Carl Roth), the membrane was incubated overnight at 4°C with a SAS-6 antibody. Secondary antibodies coupled to horseradish peroxidase were used for enhanced chemiluminescence signal detection with ECL Prime Western Blotting System (GE Healthcare).

### Image analysis

For signal quantification using ImageJ (NIH), the signal intensity from the nuclear area, as determined by DAPI staining, was normalized to DAPI intensity. The fold change was defined as a ratio between normalized signal intensity to the mean signal intensity of the WT from all replicates. The percentage of cells with centrosomes in embryos and mESCs was defined as a ratio between centrosome number manually quantified using ImageJ and the number of nuclei quantified using the IMARIS software (Bitplane; Belfast, United Kingdom). Centriole length was measured using ImageJ from nearly parallel-oriented centriole walls stained for TUB. The length was corrected for the expansion factor obtained from dividing the gel size after expansion by the size of a cover slip used for gelation (12mm). Longer-than-average centrioles were defined as centrioles longer than the sum of the mean plus one standard deviation of WT centrioles.

### Statistical analysis

To identify statistical differences between two or more groups, two-tailed Student’s t-test or one-way ANOVA with Tukey’s multiple comparisons was performed. P < 0.05 was used as the cutoff for significance. The statistical analyses were performed using Microsoft Excel (Microsoft Corporation, Redmond, WA, USA) or Prism (Graphpad; San Diego, CA, USA) and the graphs were generated using Prism.

## Data Availability

This study includes no data deposited in external repositories. Information, detailed protocols and reagents are available from the corresponding author upon reasonable request.

## Acknowledgements

We thank the CECAD *in vivo* research facility (Branko Zevnik) for generating and maintaining our mouse lines and the CECAD imaging facility for microscopy support. We thank Charlotte Meyer-Gerards for assistance with generating the centriole diagrams. We are grateful to members of the lab and colleagues for critical reading of the manuscript. The work was funded by the Deutsche Forschungsgemeinschaft (DFG, German Research Foundation) - Project-ID 73111208 - SFB829 “Molecular Mechanisms regulating Skin Homeostasis”, subproject A12 to H.B. The funders had no role in study design, data collection and analysis, decision to publish, or preparation of the manuscript.

## Author contributions

Conceptualization: M.G. and H.B.; Methodology: M.G.; Software: M.G.; Formal Analysis: M.G.; Investigation: M.G. and H.B.; Writing: M.G. and H.B.; Visualization: M.G.; Supervision, Project administration and Funding Acquisition: H.B.

## Supplementary Figure Legends

**FIG. S1.**
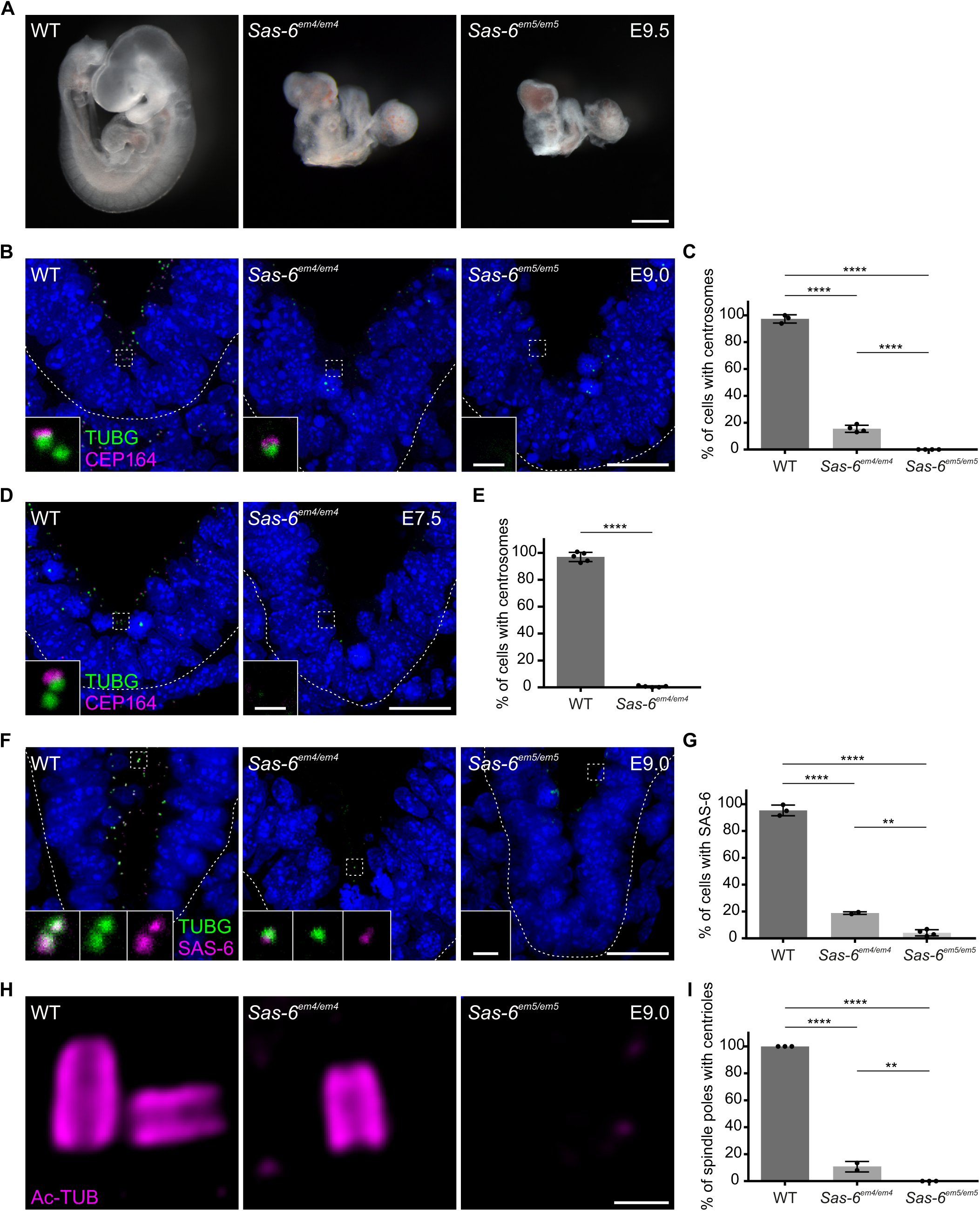
*Sas-6*^*em5/em5*^ null embryos, but not *Sas-6*^*em4/em4*^, lack centrioles. A. WT, *Sas-6*^*em4/em4*^ and *Sas-6*^*em5/em5*^ embryos at E9.5. At least five embryos were analyzed per genotype. Scale bar = 500 µm. B. Immunostaining for TUBG and CEP164 on transverse sections of WT, *Sas-6*^*em4/em4*^ and *Sas-6*^*em5/em5*^ embryos at E9.0. The sections shown are similar to those described in Fig. 1 B. Insets are magnifications of the dashed squares. Scale bars = 20 µm and 1 µm (insets). C. Quantification of the percentage of cells with centrosomes (TUBG and CEP164) in (B). Error bars represent mean ± s.d. WT: 97 ± 0.03% (n = 2843 cells from three embryos); *Sas-6*^*em4/em4*^: 16 ± 0.2% (n = 2375 from four embryos); *Sas-6*^*em5/em5*^: 0 ± 0% (n = 2152 from four embryos). ****P < 0.0001 (one-way ANOVA with Tukey’s multiple comparisons). D. Immunostaining for TUBG and CEP164 on transverse sections of WT and *Sas-6*^*em4/em4*^, embryos at E7.5. The sections shown are similar to those in Fig 1B but at an earlier stage. Insets are magnifications of the dashed squares. Scale bars = 20 µm and 1 µm (insets). E. Quantification of the percentage of cells with centrosomes (TUBG and CEP164) from (D). Five embryos per genotype were used for the quantifications. Error bars represent mean ± s.d. WT: 97 ± 3% (n = 3974 cells); *Sas-6*^*em4/em4*^: 1 ± 1% (n = 2774). ****P < 0.0001 (two-tailed Student’s t-test). F. Immunostaining for TUBG and SAS-6 in WT, *Sas-6*^*em4/em4*^ and *Sas-6*^*em5/em5*^ embryos at E9.0. Insets are magnifications of the dashed squares. Scale bars = 20 µm and 1µm (insets). G. Quantification of the percentage of cells with SAS-6 signal co-localization with TUBG in (F). Error bars represent mean ± s.d. WT: 95 ± 3% (n = 1929 cells from three embryos); *Sas-6*^*em4/em4*^: 19 ± 1% (n = 542 from two embryos); *Sas-6*^*em5/em5*^: 4 ± 2% (n = 2458 from four embryos). ****P < 0.0001, **P < 0.01 (one-way ANOVA with Tukey’s multiple comparisons). H. Immunostaining for Ac-TUB on U-ExM sections from E9.0 embryos. Scale bar = 200 nm. I. Quantification of the percentage of spindle poles with centrioles in (H). Error bars represent mean ± s.d. WT: 100 ± 0% (n = 65 spindle poles from three embryos); *Sas-6*^*em4/em4*^: 11 ± 0.03% (n = 62 from two embryos); *Sas-6*^*em5/em5*^: 0 ± 0% (n = 45 from three embryos). ****P < 0.0001, **P < 0.01 (one-way ANOVA with Tukey’s multiple comparisons).

**FIG. S2.**
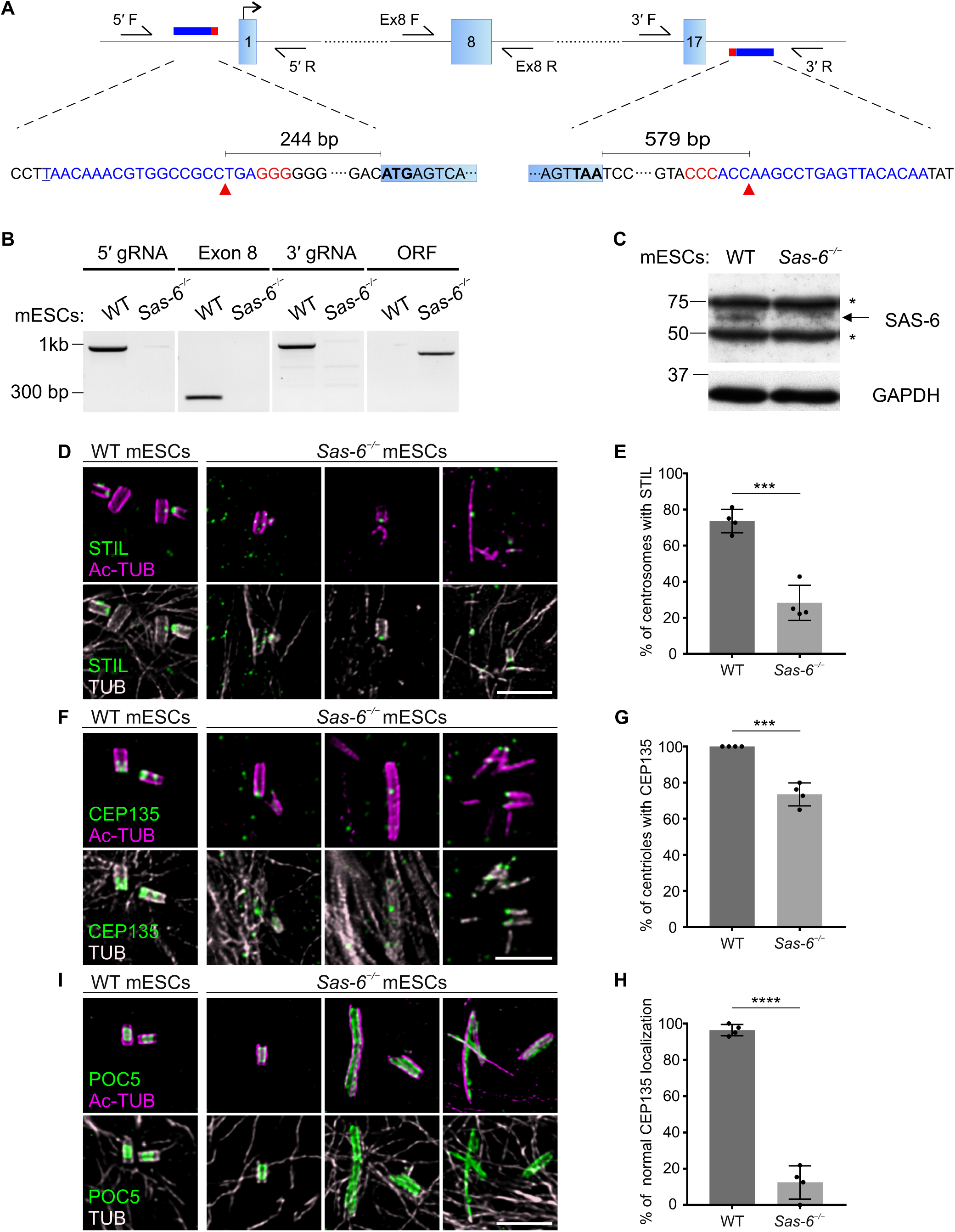
Deletion of *Sas-6* ORF in mESCs and analyzing centriole integrity. A. Schematic showing the CRISPR/Cas9 strategy to delete the *Sas-6* ORF. Exons are represented by blue boxes, gRNAs by blue horizontal lines, PAM sites in red and Cas9 cut sites by red arrowheads. The sequences complementary to the gRNAs are in blue letters and the PAM sequences in red letters. Half arrows indicate the primers used for PCR analyses in (B). B. Confirmation of the *Sas-6* deletion in *Sas-6*^*−/−*^ mESCs by genomic PCR. The agarose gel shows the amplified products using the primers depicted in (A) from the region flanking the 5′ gRNA (5′ F and 5′ R, band = 977 bp), Exon 8 (Ex8 F and Ex8 R, band = 281 bp), 3′ gRNA (3′ F and 3′ R, band = 992 bp), *Sas-6* ORF (5′ F and 3′ R, 825 bp in *Sas-6*^*−/−*^, 34,349 bp in WT, product too long to be amplified). C. Western blot analysis of SAS-6 in WT and *Sas-6*^*−/−*^ mESCs extracts. Asterisks mark non-specific bands. D. Immunostaining for Ac-TUB and STIL of U-ExM expanded centrioles from WT and *Sas-6*^*−/−*^ mESCs. E. Quantification of the percentage of centrosomes with STIL in (D) from four independent experiments. Error bars represent mean ± s.d. WT: 74 ± 6% (n = 72 centrosomes); *Sas-6*^*−/−*^ : 29 ± 8% (n = 94). ***P < 0.001, (two-tailed Student’s t-test). Scale bar = 1 µm. F. Immunostaining for Ac-TUB and cartwheel protein (CEP135) of U-ExM expanded centrioles from WT and *Sas-6*^*−/−*^ mESCs. Scale bars = 1 µm. G. Quantifications of the percentage of centrioles with associated CEP135 in (F) from four independent experiments. Error bars represent mean ± s.d. WT: 100 ± 0% (n = 160 centrioles); *Sas-6*^*−/−*^ : 73 ± 6% (n = 98). ***P < 0.001, (two-tailed Student’s t-test). H. Quantifications of normal or WT-like localization of CEP135 in (F) from four independent experiments. Error bars represent mean ± s.d. WT: 96 ± 3% (n = 160 centrioles); *Sas-6*^*−/−*^ : 12 ± 8% (n = 98). ****P < 0.0001, (two-tailed Student’s t-test). I. Immunostaining for Ac-TUB and inner scaffold protein (POC5) of U-ExM expanded centrioles from WT and *Sas-6*^*−/−*^ mESCs. Scale bar = 1 µm.

**Table S1.**
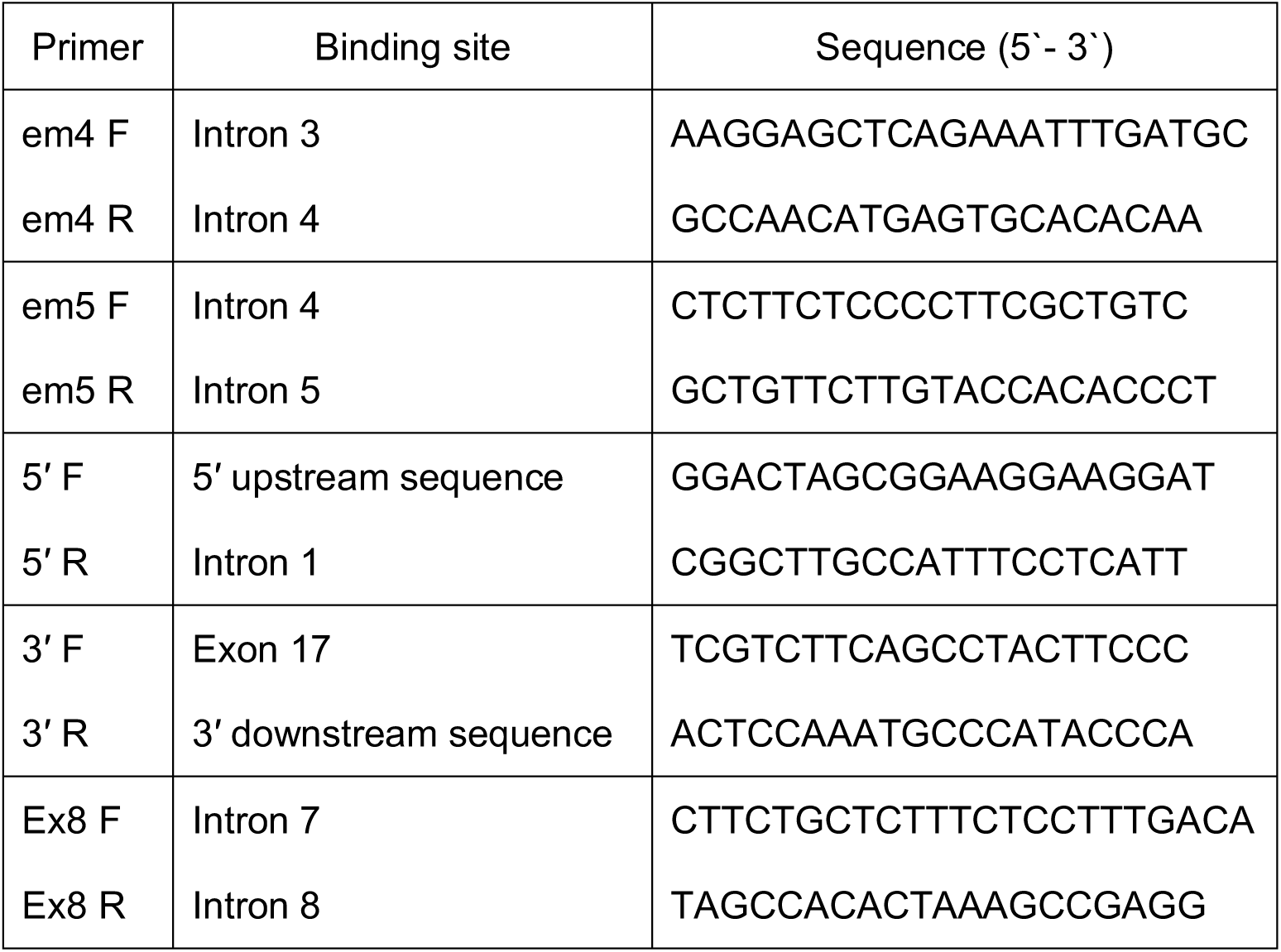
List of used primers in the mouse *Sas-6* locus.

## References

Bangs FK, Schrode N, Hadjantonakis AK, Anderson KV (2015) Lineage specificity of primary cilia in the mouse embryo. Nat Cell Biol 17: 113–122

Bayless BA, Giddings TH, Jr., Winey M, Pearson CG (2012) Bld10/Cep135 stabilizes basal bodies to resist cilia-generated forces. Mol Biol Cell 23: 4820–4832

Bazzi H, Anderson KV (2014a) Acentriolar mitosis activates a p53-dependent apoptosis pathway in the mouse embryo. Proc Natl Acad Sci U S A 111: E1491–1500

Bazzi H, Anderson KV (2014b) Centrioles in the mouse: cilia and beyond. Cell Cycle 13: 2809

Bond J, Roberts E, Springell K, Lizarraga S, Scott S, Higgins J, Hampshire DJ, Morrison EE, Leal GF, Silva EO et al (2005) A centrosomal mechanism involving CDK5RAP2 and CENPJ controls brain size. Nature Genetics 37: 353–355

Bornens M (2012) The centrosome in cells and organisms. Science 335: 422–426

Bryja V, Bonilla S, Arenas E (2006) Derivation of mouse embryonic stem cells. Nat Protoc 1: 2082–2087

Chu VT, Weber T, Graf R, Sommermann T, Petsch K, Sack U, Volchkov P, Rajewsky K, Kuhn R (2016) Efficient generation of Rosa26 knock-in mice using CRISPR/Cas9 in C57BL/6 zygotes. BMC Biotechnol 16: 4

Conduit PT, Wainman A, Raff JW (2015) Centrosome function and assembly in animal cells. Nat Rev Mol Cell Biol 16: 611–624

Courtois A, Schuh M, Ellenberg J, Hiiragi T (2012) The transition from meiotic to mitotic spindle assembly is gradual during early mammalian development. J Cell Biol 198: 357–370

Gambarotto D, Zwettler FU, Le Guennec M, Schmidt-Cernohorska M, Fortun D, Borgers S, Heine J, Schloetel JG, Reuss M, Unser M et al (2019) Imaging cellular ultrastructures using expansion microscopy (U-ExM). Nat Methods 16: 71–74

Gueth-Hallonet C, Antony C, Aghion J, Santa-Maria A, Lajoie-Mazenc I, Wright M, Maro B (1993) gamma-Tubulin is present in acentriolar MTOCs during early mouse development. Journal of Cell Science 105 (Pt 1): 157–166

Gupta H, Rajeev R, Sasmal R, Radhakrishnan RM, Anand U, Chandran H, Aparna NR, Agasti S, Manna TK (2020) SAS-6 Association with gamma-Tubulin Ring Complex Is Required for Centriole Duplication in Human Cells. Curr Biol 30: 2395–2403 e2394

Hafner M, Wenk J, Nenci A, Pasparakis M, Scharffetter-Kochanek K, Smyth N, Peters T, Kess D, Holtkotter O, Shephard P et al (2004) Keratin 14 Cre transgenic mice authenticate keratin 14 as an oocyte-expressed protein. Genesis 38: 176–181

Higginbotham H, Bielas S, Tanaka T, Gleeson JG (2004) Transgenic mouse line with green-fluorescent protein-labeled Centrin 2 allows visualization of the centrosome in living cells. Transgenic research 13: 155–164

Hilbert M, Erat MC, Hachet V, Guichard P, Blank ID, Fluckiger I, Slater L, Lowe ED, Hatzopoulos GN, Steinmetz MO et al (2013) Caenorhabditis elegans centriolar protein SAS-6 forms a spiral that is consistent with imparting a ninefold symmetry. Proc Natl Acad Sci U S A 110: 11373–11378

Kantsadi AL, Hatzopoulos GN, Gonczy P, Vakonakis I (2022) Structures of SAS-6 coiled coil hold implications for the polarity of the centriolar cartwheel. Structure 30: 671–684 e675

Khan MA, Rupp VM, Orpinell M, Hussain MS, Altmuller J, Steinmetz MO, Enzinger C, Thiele H, Hohne W, Nurnberg G et al (2014) A missense mutation in the PISA domain of HsSAS-6 causes autosomal recessive primary microcephaly in a large consanguineous Pakistani family. Hum Mol Genet 23: 5940–5949

Kitagawa D, Vakonakis I, Olieric N, Hilbert M, Keller D, Olieric V, Bortfeld M, Erat MC, Fluckiger I, Gonczy P et al (2011) Structural basis of the 9-fold symmetry of centrioles. Cell 144: 364–375

Kratz AS, Barenz F, Richter KT, Hoffmann I (2015) Plk4-dependent phosphorylation of STIL is required for centriole duplication. Biology open 4: 370–377

Lambrus BG, Holland AJ (2017) A New Mode of Mitotic Surveillance. Trends Cell Biol 27: 314–321

Leidel S, Delattre M, Cerutti L, Baumer K, Gonczy P (2005) SAS-6 defines a protein family required for centrosome duplication in C. elegans and in human cells. Nat Cell Biol 7: 115–125

Lin YC, Chang CW, Hsu WB, Tang CJ, Lin YN, Chou EJ, Wu CT, Tang TK (2013) Human microcephaly protein CEP135 binds to hSAS-6 and CPAP, and is required for centriole assembly. EMBO J 32: 1141–1154

Mahmood T, Yang PC (2012) Western blot: technique, theory, and trouble shooting. N Am J Med Sci 4: 429–434

Marino S, Vooijs M, van Der Gulden H, Jonkers J, Berns A (2000) Induction of medulloblastomas in p53-null mutant mice by somatic inactivation of Rb in the external granular layer cells of the cerebellum. Genes Dev 14: 994–1004

Marshall WF (2007) What is the function of centrioles? J Cell Biochem 100: 916–922

Nabais C, Peneda C, Bettencourt-Dias M (2020) Evolution of centriole assembly. Curr Biol 30: R494–R502

Nakazawa Y, Hiraki M, Kamiya R, Hirono M (2007) SAS-6 is a cartwheel protein that establishes the 9-fold symmetry of the centriole. Curr Biol 17: 2169–2174

Nigg EA, Holland AJ (2018) Once and only once: mechanisms of centriole duplication and their deregulation in disease. Nat Rev Mol Cell Biol 19: 297–312

Qiao R, Cabral G, Lettman MM, Dammermann A, Dong G (2012) SAS-6 coiled-coil structure and interaction with SAS-5 suggest a regulatory mechanism in C. elegans centriole assembly. EMBO J 31: 4334–4347

Rodrigues-Martins A, Bettencourt-Dias M, Riparbelli M, Ferreira C, Ferreira I, Callaini G, Glover DM (2007) DSAS-6 organizes a tube-like centriole precursor, and its absence suggests modularity in centriole assembly. Curr Biol 17: 1465–1472

Tang C-JC, Lin S-Y, Hsu W-B, Lin Y-N, Wu C-T, Lin Y-C, Chang C-W, Wu K-S, Tang TK (2011) The human microcephaly protein STIL interacts with CPAP and is required for procentriole formation. The EMBO journal 30: 4790–4804

Troder SE, Ebert LK, Butt L, Assenmacher S, Schermer B, Zevnik B (2018) An optimized electroporation approach for efficient CRISPR/Cas9 genome editing in murine zygotes. PLoS One 13: e0196891

van Breugel M, Hirono M, Andreeva A, Yanagisawa HA, Yamaguchi S, Nakazawa Y, Morgner N, Petrovich M, Ebong IO, Robinson CV et al (2011) Structures of SAS-6 suggest its organization in centrioles. Science 331: 1196–1199

Vulprecht J, David A, Tibelius A, Castiel A, Konotop G, Liu F, Bestvater F, Raab MS, Zentgraf H, Izraeli S et al (2012) STIL is required for centriole duplication in human cells. Journal of Cell Science 125: 1353–1362

Wang WJ, Acehan D, Kao CH, Jane WN, Uryu K, Tsou MF (2015) De novo centriole formation in human cells is error-prone and does not require SAS-6 self-assembly. Elife 4

Wang X, Yang P (2008) In vitro differentiation of mouse embryonic stem (mES) cells using the hanging drop method. J Vis Exp

Xiao C, Grzonka M, Meyer-Gerards C, Mack M, Figge R, Bazzi H (2021) Gradual centriole maturation associates with the mitotic surveillance pathway in mouse development. EMBO Rep 22: e51127

Yoshiba S, Tsuchiya Y, Ohta M, Gupta A, Shiratsuchi G, Nozaki Y, Ashikawa T, Fujiwara T, Natsume T, Kanemaki MT et al (2019) HsSAS-6-dependent cartwheel assembly ensures stabilization of centriole intermediates. Journal of cell science 132

